# Long-term laboratory Drosophila populations prefer ancestral nutritional cues from the environment

**DOI:** 10.1101/2025.04.17.649384

**Authors:** Pedro Simões, Marta A. Antunes, Margarida Matos

**Affiliations:** CE3C – Centre for Ecology, Evolution and Environmental Changes & CHANGE – Global Change and Sustainability Institute, Lisboa, Portugal; Departamento de Biologia Animal, Faculdade de Ciências, Universidade de Lisboa, Lisboa, Portugal

**Author notes:** corresponding author, **email:**. Departamento de Biologia Animal, C2 building, Lab 2.1.40, Faculdade de Ciências, Universidade de Lisboa, Lisboa, Portugal. MAA -, MM –.

**Keywords:** Plasticity, Drosophila, Ecological niche, Fecundity, Fruit host

## Abstract

Plasticity can help populations cope with environmental changes namely by exploring different ecological niches. Addressing plasticity for nutritional responses in a range of fruit hosts potentially used by Drosophila may be essential in predicting the capacity of insects to colonize new environments or return to ancestral ones. Here we test for differences in oviposition performance, reproductive success and juvenile viability in different host fruits in the colonizing species *Drosophila subobscura* Collin, and compare them with those of the laboratory maintenance medium to which populations adapted for ∼150 generations. We question: does *Drosophila subobscura* show plasticity associated with different fruit hosts? Is performance better in the long-term maintenance (control) medium? We observed a higher fecundity, reproductive success and juvenile viability of flies maintained in the fruit media versus the control, but no differences between fruits. Our experiment shows that long-term lab populations of *Drosophila subobscura* can still assess environmental cues of new substrates allowing for flexible adaptive plasticity to occur through increased fecundity and reproductive success in fruit hosts relative to the control conditions. Importantly, this ability was not lost during long-term evolution in a benign, homogeneous environment. Furthermore, the high performance across fruits reinforces its status as a generalist species and further attests its potential to colonize different ecological settings.

## Introduction

Insects are currently declining worldwide as a result of exposure to rapidly deteriorating environments (Wagner et al. 2021; Harvey et al. 2023). In this context, the plastic ability of insect species to show high levels of reproductive success when exposed to different ecological niches is a clear advantage (Chevin et al. 2010; Beldade et al. 2011; Raubenheimer et al. 2012). A relevant ecological feature of insects – including Drosophila - is the potential to select adequate substrates (i.e. trophic niches) for feeding and oviposition (Markow and O’Grady 2005, 2008; Raubenheimer et al. 2012). Oviposition performance – that encompasses both the ability to cognitively decide whether a site is suitable for oviposition and the capacity to lay an adequate number of eggs – is a major factor that can drive the expansion of insects in new environments (Foucaud et al. 2016; Fanara et al. 2023). The decision of insects to oviposit in a particular site/ host is shaped by the integration of a variety of “external” environmental cues that potentially reflect host nutrient content and chemistry (mechanosensory, chemosensory and visual) as well as intrinsic factors of the organism (e.g., its physiological status) – see (Yang et al. 2008; Joseph et al. 2009; Joseph and Carlson 2015; Karageorgi et al. 2017; Zhang et al. 2020; Fanara et al. 2023). Ultimately, the ability of organisms to thrive in different hosts will depend on the availability of resources / nutrients on their diet – likely with a prominent role of proteins - that will allow for egg production as well as their successful development (Mirth et al. 2019; Shu et al. 2022; Poças et al. 2022); see also references below.

Studies on oviposition performance and reproductive success in different trophic niches are relevant to understand the ability of organisms to cope with specific hosts and determine the invasive capacity of different species. Drosophila is an excellent species for such studies considering its ability to explore a wide range of ecological resources, namely using different fruits as hosts for feeding and oviposition (Mueller 1985; Markow and O’Grady 2005, 2008; Raubenheimer et al. 2012). Attractiveness to fruit hosts in Drosophila can depend on several aspects, namely the role of volatile chemical compounds present in different fruits as well as other cues, e.g. associated with the taste, colour and stiffness of the substrate that can provide varied information on the nutritional status of the host fruit (Abraham et al. 2015; Karageorgi et al. 2017; Zhang et al. 2020).

Nutritional geometry addresses how organisms respond to changes in host nutrient availability (Simpson and Raubenheimer, 1993). One key aspect of nutritional geometry of hosts is the protein to carbohydrate (P:C) ratio (Lee et al., 2008; Sario et al., 2022; Young et al., 2018) with higher P:C ratios more frequently associated with increased fecundity but also decreased lifespan in fruit flies (Lee et al. 2008; Fanson et al. 2012; Jang and Lee 2018; Shu et al. 2022), but see (Silva-Soares et al. 2017; Young et al. 2018) for different results in *D. suzukii*). Increased fecundity is likely due to the positive effects of dietary protein on oogenesis (Jang and Lee 2018; Mirth et al. 2019)and also the positive impact of increased body size (Jang and Lee 2018; Poças et al. 2022). Higher P:C ratios have also been related to increased larval development (Shu et al. 2022; Poças et al. 2022). On the other hand, increased values of carbohydrates - a source of energy for the organism - can lead to higher starvation resistance (Shu et al. 2022) and lifespan (Lee et al. 2008; Fanson et al. 2012)up to a given threshold (Jang and Lee, 2018).

Studies measuring performance under different fruits have been mostly done on *Drosophila suzukii* (Silva-Soares et al. 2017; Young et al. 2018; Olazcuaga et al. 2019; Sario et al. 2022), due to its fruit pest status (Asplen et al. 2015). However, *D. suzukii* might not be the best representative of the Drosophila genus in terms of habitat selection, considering its singular attraction to ripe fruit (contrary to most Drosophila species, that lay eggs on overripe fruit). In fact, in a study of nutritional geometry in *D. suzukii* and *D. biarmipes*, Silva-Soares et al. (2017) showed that *D. suzukii* is able to colonize a wider range of food substrates than *D. biarmipes* highlighting the role of nutritional performance and feeding behaviour in the process of host colonization, coupled with morphological (e.g. serrated ovipositor) and sensory changes in this species (see also (Karageorgi et al. 2017). It is important to analyse how responses in a range of fruit hosts vary in other generalist Drosophila species other than *D. suzukii*, namely those with distinct habitat (feeding and breeding) preferences (e.g. that use more overripe, rotting fruit as substrate). Addressing the amplitude and host types used by different species may be an essential factor in predicting their capacity to colonize and thrive in new environments.

*Drosophila suboboscura* is a case study of a successful colonizing species, having quickly spread through South and North America from its European origin (Rezende et al. 2010). It is a generalist species with a saprophytic nutrition mostly based on decaying plant material and fruits, as well as yeast and microbes (Markow and O’Grady 2008). Despite being an emblematic example of a successfully colonizing species, several aspects of *D. subobscura* ecology, including its ability to cope with distinct nutritional environments, are surprisingly unknown. A study on oviposition performance using introduced and native populations of this species has shown that differences between populations rely on fecundity (“quantity” of oviposition) rather than on learning (“quality” of oviposition, based on oviposition site choice) - (Foucaud et al. 2016). In this study flies were exposed to media with different fruit odors (e.g. banana or strawberry) and did not focus on the oviposition in the fruits themselves or at least in media including fruit nutrients. This is a relevant missing experiment for a better understanding of the ability of this species to cope with new environments. Increased ability to explore different trophic niches may be an important advantage for this species in face of new and potentially deteriorating environments.

In this study, we will directly test for differences in *D. subobscura* oviposition performance (i.e. fecundity), reproductive success (i.e. number of offspring) and juvenile viability in different hosts. We will use populations already adapted to the laboratory conditions for many generations. These populations showed marked temporal increases in their reproductive performance during the first ∼30 generations of evolution in the new laboratorial environment (Simões et al., 2017). For new hosts, we chose four fruits that vary in their nutritional geometry (levels of protein and carbohydrates) with described effects on Drosophila key life history traits (mostly on *D. suzukii*: see (Abraham et al. 2015; Olazcuaga et al. 2019; Shu et al. 2022; Sario et al. 2022)). The goal is to compare the performance of the females (in egg laying and reproductive success) among these fruits as well as between them and the standard laboratory maintenance (control) medium (to which populations are already adapted).

Specifically, we question:

1. Does *Drosophila subobscura* show plasticity in fecundity and/or reproductive success associated with the different fruit hosts? Is it able to lay eggs and develop in a range of different hosts with varying nutritional geometry?
2. Do *D. subobscura* lab populations perform better in the nutritional diet they have been continuously exposed to for almost 150 generations? Or are they able to adjust their performance to better explore new hosts?

Our general expectation is that populations will perform better in the culture medium they adapted to for many generations. This adaptation to the standard maintenance medium can potentially have involved the ability to recognize specific olfactory, gustatory and tactile (e.g. roughness and hardness) of such medium as “positive”, while potentially reducing its perception and/or attractiveness to less frequently experienced cues particularly if involving costly mechanisms (Murren et al., 2015; Snell-Rood et al., 2018). However, it is also possible that, on the contrary, volatiles and gustatory cues (e.g. due to increased sugar content) in some fruits are sufficiently attractive, leading to an increased fecundity in such media relative to the control medium. It is also an open question whether increased sugar levels in fruits will directly allow for a higher fecundity (due to improved oogenesis) as well as a higher juvenile viability and hence higher reproductive success relative to the control media. Importantly, the standard medium does not include simple sugars (see Material and Methods), which may be an important nutritional source. When comparing the different fruit media, we expect that fecundity and developmental success, and ultimately reproductive success will be enhanced in media with higher levels of protein relative to carbohydrates (high P:C ratios) considering the relevance of protein content for reproduction.

## Material and Methods

*Drosophila subobscura* populations used in this experiment – the PT populations – were derived from a natural collection in Adraga (Portugal) that already evolved in the laboratory environment for 145 generations when the assay was done. The initial outbred population was three-fold replicated few generations after collection, generating three replicate populations (labelled PT1-3). Populations were maintained in a standard medium (hereafter called ‘control medium’), in 28-day cycle at 18°C and 12L:12D photoperiod, with control adult and egg densities - see details in (Simões et al. 2017).

We performed an experimental assay to test how *Drosophila subobscura* adults differed in egg laying and juvenile viability when exposed to different fruit media as well as the control medium. This assay involved 5 different media: the control medium and four new media that included fruit juices of raspberry, blueberry, strawberry, andwhite grapes. . These fruits involved a varying nutritional status based on information from other studies (namely different Protein: Carbohydrate ratios – P:C, see (Olazcuaga et al. 2019; Shu et al. 2022; Sario et al. 2022) as well as their natural occurrence in Portugal. Furthermore, there is no evidence that *D. subobscura*, a generalist species, avoids breeding or feeding in any of the chosen fruits. The control medium used the following ingredients: Brewer’s dead yeast (28.75 g), corn flour (27.5 g), agar (3,75 g) and nipagin (3 g) in a total of 325 ml of medium. Fruit media used the same ingredients, but corn flour was replaced by fruit purée (160g) for a final volume of 385 ml. Commercially available frozen fruits were used to control for the ripeness of the different fruits. Fruit media used a slightly higher total volume (+50 ml) than the control medium to allow for all ingredients to be successfully dissolved (reducing the excess of thickness in the medium). This also allowed to minimize differences in medium thickness while maintaining approximate concentrations of the ingredients across media.

The nutritional information available for the fruits used in this experiment allowed us to calculate the different protein to carbohydrate (P:C) and protein to total sugar (P:S) ratios for each fruit (see Table 1). P:C and P:S ratios for each fruit were very similar reflecting that most of the carbohydrate content of fruits was due to simple sugars. P:S values are quite comparable to the ones reported by Shu et al. (2022) for the same fruits. In contrast, the control medium had clearly discrepant values of P:C and P:S ratio with the latter much higher than the former, indicating that most carbohydrate content (provided by corn flour, see above) were not simple sugars.

**Table 1.**
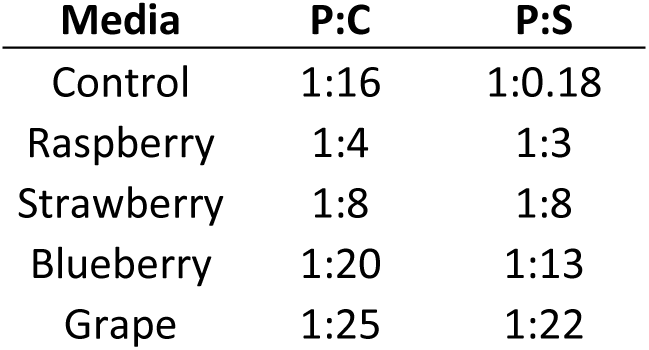
Nutritional information - Protein to Carbohydrate (P:C) and Protein to Sugar (P:S) ratios - for each culture media.

| **Media** | **P:C** | **P:S** |
| --- | --- | --- |
| Control | 1:16 | 1:0.18 |
| Raspberry | 1:4 | 1:3 |
| Strawberry | 1:8 | 1:8 |
| Blueberry | 1:20 | 1:13 |
| Grape | 1:25 | 1:22 |

The assay involved 16 pairs (of one virgin female and one virgin male per vial) per population and fruit media, in a total of 240 vials (16 pairs x 3 populations x 5 media). To obtain individuals for the assay, 18 vials with 70 eggs each were collected for each replicate population and allowed to develop in the control medium. Assayed individuals were all obtained in the 20^th^ day since egg collection, with the pairing for the assay done around 2 to 3 hours post emergence. The assay was performed in blocks, with each “block” corresponding to vials from one replicate population that were assayed in the same experimental rack (e.g. Block 1 includes all vials from PT1 population). The three blocks were assayed in synchrony. Pairs of mated flies were transferred to fresh media every day and the presence of eggs was checked daily. Eggs were counted on days 6 and 7 of the imagos’ lives. At the last day of the assay, flies were removed from vials and the eggs laid in the last 24 hours were allowed to develop. We chose days 6 and 7 as they are within the age interval of flies that contribute to the next generation in our maintenance system, expectedly when the strength of selection is higher. The reproductive success of each pair of flies was assessed by counting the number of emerging offspring (for a total of 10 days since first emergence). Several traits were measured in this assay: Age of first reproduction (i.e. number of days until the first egg-laying); Total Fecundity on days 6 and 7 (as a measure of fecundity close to maturity); Fecundity on day 7 (F7, the last day of the assay, used to estimate reproductive success); Reproductive success (number of offspring derived from F7) and Juvenile Viability (ratio between reproductive success and F7) – raw data is available in Tables S1-S2.

We aim to test whether fecundity, reproductive success and Viability (the dependent variables) vary as a function of exposure to different fruit media during the adult stage as well as during development of the offspring (for reproductive success and juvenile viability). Fecundity (Total Fecundity and F7) and reproductive success were analysed with Generalized linear mixed-effects models (GLMM, glmmTMB command) on the individual data (each mating pair) and two distributions for count data were tested - quasipoisson and negative binomial. We chose the model with quasipoisson distribution, based on its lowest values of Akaike information criterion (AIC). “Sum to zero” contrasts were applied for each factor. Significance levels were obtained by applying Type III Wald chisquare tests. Viability was analysed with a linear mixed-effects model with the viability data from each individual being arcsine-transformed to better fit normality. To avoid unreliable estimates of viability due to low egg number, vials with F7 values below 5 were not considered in the analysis of viability. Age of first reproduction was also analysed with a linear mixed-effects model with Gaussian distribution due to convergence problems when applying the GLMM models – see data in Supplementary Material.

The model applied for all traits was the following:

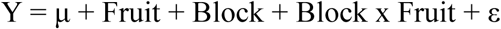

In the model Block is a random factor that accounts for the variation between sets of racks, with each block corresponding to a given replicate population assayed in the same set of experimental racks (containing vials from all 5 media) with pseudo-randomization of vials. Y is the studied trait (age of first reproduction, fecundity, reproductive success or viability). Fruit is the fixed factor corresponding to the 5 different media (the control medium and the 4 different fruit media). The interaction Block x Fruit corresponds to the variation between replicate populations after accounting for the block effect (conceptually corresponding to the error level of the fixed factor). To specifically test for differences between media using different fruits a similar model as described above was applied but excluding data from the control medium. Tukey post-hoc tests were applied to test for differences between all possible combinations (pairs) of culture media and a contrast was also performed between all fruit media vs control for each trait.

Statistical analyses and graphs were performed with R v4.3.3, using lme4, glmmTMB, car, emmeans and ggplot2 packages (Wickham 2016; Brooks et al. 2017; Fox and Weisberg 2019).

## Results

We observed significant overall differences between culture media for both total fecundity (F6-7) and reproductive success (see Table 2, Figures 1 and 2). When dissecting differences between all fruit media vs control, we observed a significantly higher performance in fruit media relative to the control for both traits (total fecundity: Z ratio = 3.595, d.f. = inf, p = 0.0003, corresponding to a 26.6 % increase in average performance in fruit media vs control; reproductive success: Z ratio = 3.030, d.f. = inf, p = 0.002, corresponding to a 28.6 % increase in the average performance in fruit media vs control). We further analysed variation between media by applying pairwise comparisons. We observed significantly higher fecundity in the raspberry, blueberry and grape media relative to the control medium; and higher reproductive success in the raspberry and blueberry vs the control medium (Table S3). When excluding the control medium, no significant differences were observed among media with different fruits (Table S4). While raspberry was the fruit host that consistently showed higher average total fecundity and reproductive success, variation between replicate populations was very high (see Figures 1 and 2).

**Figure 1.**
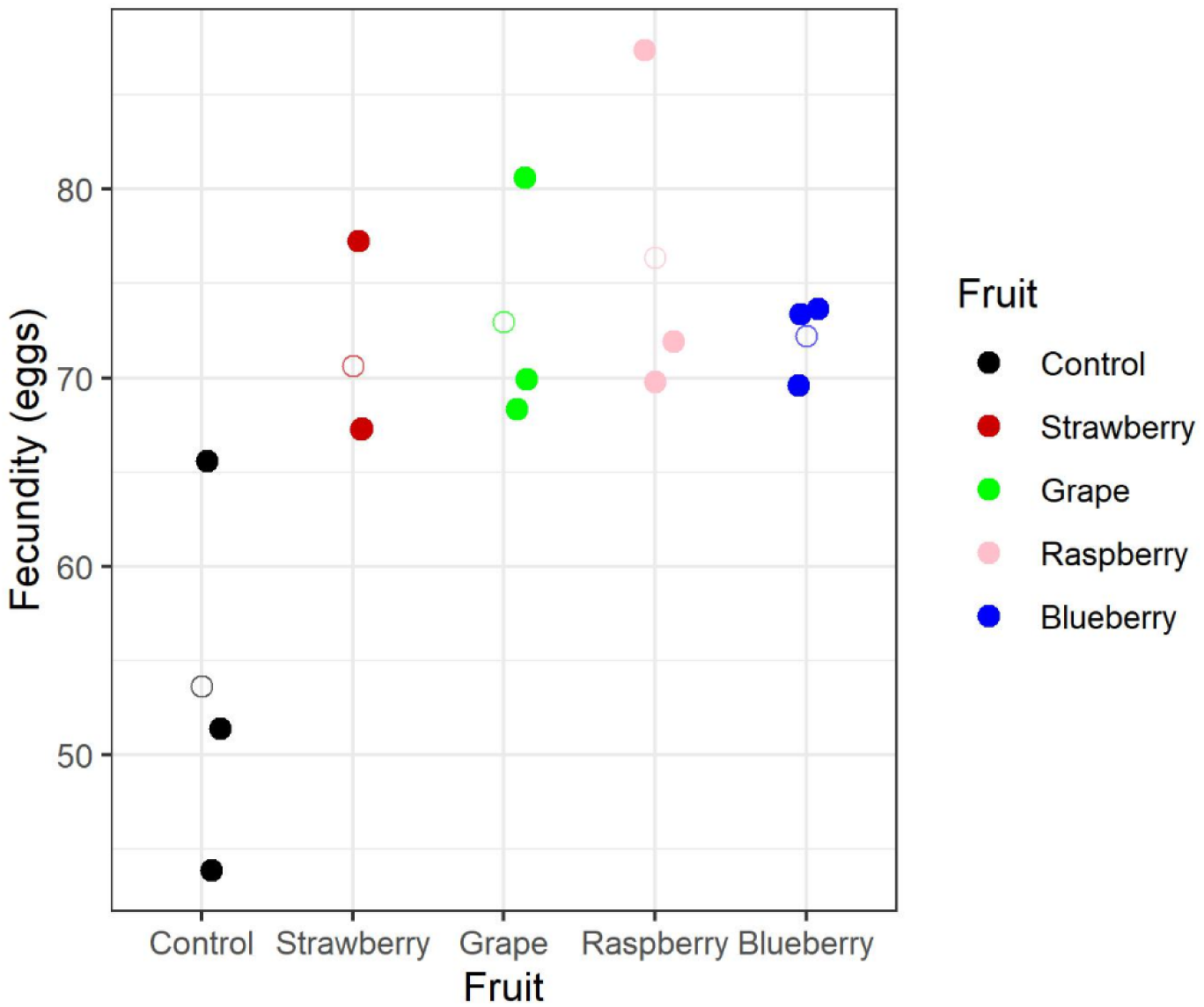
Total Fecundity across different media. Full circles represent each replicate population analysed in the study, while the open circles represent the average value of the three replicate populations.

**Figure 2.**
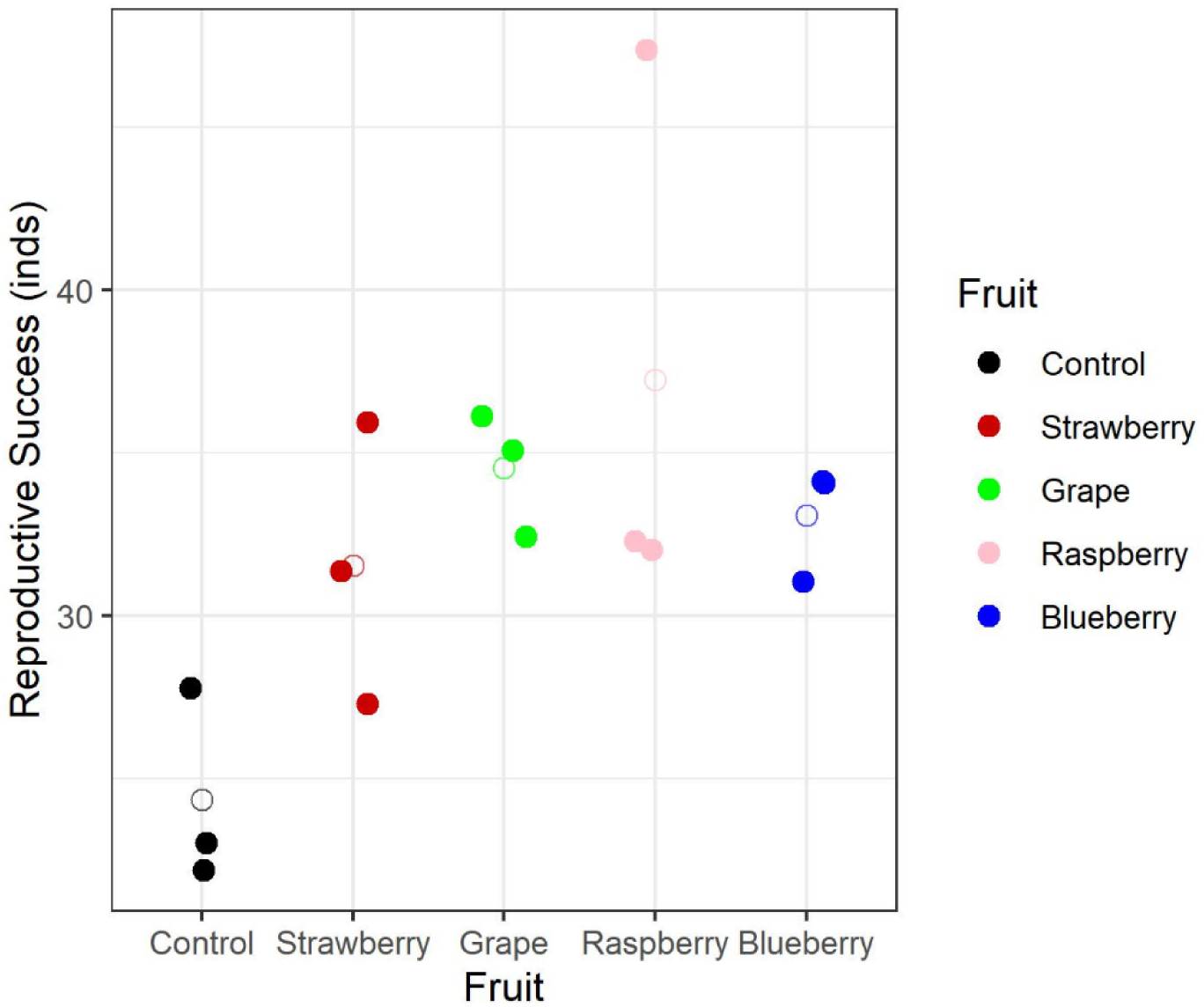
Reproductive Success across different media. Full circles represent each replicate population analysed in the study, while the open circles represent the average value of the three replicate populations.

**Table 2.** Overall differences in performance between culture media for each trait analysed.

| Trait | d.f. | $\chi^2$ / F | p-value |
| --- | --- | --- | --- |
| **Total Fecundity (F6-7)** | **4** | **13.987** | **0.007** |
| **Reproductive success** | **4** | **11.499** | **0.021** |
| Fecundity day 7 (F7) | 4 | 6.171 | 0.187 |
| Age of First Reproduction (A1R) | 4, 7.980 | 2.067 | 0.178 |
| Viability | 4, 7.94 | 2.431 | 0.133 |

On the other hand, the different media had no overall significant effect on other traits, namely fecundity of the last day of the assay (F7), age of first reproduction or juvenile viability (Table 2 and Figures S1-S3). However, for both F7 and viability, performance in fruit media was significantly better than the control (Z ratio = 2.123, d.f. = inf, p = 0.034 for F7; Z ratio = 2.682, d.f. = 8.35, p = 0.027 for viability) – see also Figures S1 and S3. In the case of viability, this difference was likely due to a higher performance in raspberry, with marginally significant differences relative to the control medium (Table S3). As described above for fecundity and reproductive success, no differences were found between media with different fruits for these traits (analysis excluding the control, see Table S4).

## Discussion

In this experiment we assessed the plastic response of long-term lab adapted *Drosophila subobscura* populations subjected to different feeding and breeding environments, including several fruit hosts and the control medium (used in their long-term maintenance). We found a common trend of clearly higher fecundity and reproductive success when *D. subobscura* were maintained in fruit hosts rather than in their control medium, but no differences in performance between the different fruits.

It is revealing that we observe increases in key life-history traits when *D. subobscura* flies are subjected to fruit hosts compared to the environment in which they have evolved for almost 150 generations, to which they have already adapted (Simões et al. 2017). Although we did not measure plasticity at the inception of lab evolution it is very unlikely that we would have found an initial better performance in the control medium relative to fruit hosts (i.e. a change in plasticity sign), considering the initially quite low reproductive performance of recently caught populations in the new lab conditions (including the new maintenance medium) and their consistent improvement in performance during the first ∼30 generations (Simões et al. 2017). On the contrary, it seems likely that an even higher (initial) plasticity might occur in the recently introduced lab populations than that reported here, due to a better performance of flies in fruit vs control media, being the latter a quite new medium at the time. In any case, our findings suggest that these long-term lab-maintained *D. subobscura* flies are still apt to interpret environmental cues of fruits, with increased oviposition and reproductive success in a new feeding and oviposition environment. It is possible that this derives from a cognitive assessment of environmental cues, a valuable asset that was not lost even in a confined, homogeneous environment (e.g. (Murren et al., 2015; Snell-Rood et al., 2018), as is the case of the lab environment. Curiously, plasticity in response to other environments, such as new thermal challenges had already been described in these long-term laboratory populations (Fragata et al. 2016; Simões et al. 2020; Santos et al. 2021). Alternatively, the observed fecundity increases in fruit media might be directly modulated by the nutrient levels - namely simple sugars available-in the diet (Lee et al. 2008) through nutrient-sensing pathways (e.g. Kim and Neufeld 2015). This last hypothesis is supported by the fact that fecundity was measured after one week of exposure to each fruit diet.

The consistent fecundity differences observed between fruit media and the standard one indicate that these long-term lab flies can adjust their oviposition effort depending on the host media - even though each individual fly in the experiment did not have the possibility to choose between them. Interestingly, a pilot choice experiment where flies were indeed exposed simultaneously to a fruit medium vs the control medium (that we performed using the same fruit media as described here) suggests that these *D. subobscura* populations were significantly more attracted to the raspberry and blueberry media than to the control one (data not shown). These findings are in line with the fact that these two media are amongst those with higher fecundity in the present no-choice study. Our experiment does not allow to distinguish which cues (e.g. olfactory, tactile, visual) drove the performance observed. Chemosensory elements, e.g. volatile compounds and/or increased sugar content in fruit media may have played a role in the enhanced oviposition in such substrates (Abraham et al., 2015; Karageorgi et al., 2017; Yang et al., 2008).In addition, simple sugar content can promote egg-laying decision in Drosophila in some contexts, e.g. dependent on the protein:carbohydrate ratio (Yang et al., 2008). However, we cannot exclude other clues, namely visual and tactile (Karageorgi et al. 2017; Zhang et al. 2020).

While the increased oviposition (discussed above) can in part explain the higher reproductive success in fruits relative to control conditions, it is also likely that the nutritional components of the fruit media also played a role in the successful development of eggs to adults. In fact, this is corroborated by the overall increased juvenile viability of fruit media relative to the control media, though not as clear as for reproductive success. This overall high juvenile viability across different host fruits is in agreement with the species’ described potential to colonize novel environments (e.g. (Rezende et al. 2010; Foucaud et al. 2016)). The very low levels of protein in each fruit indicates that most protein content is supplied by the brewer’s yeast in the media, a major source of dietary protein, common to all media analysed in this study. In fact, inclusion of yeast in diet has been shown to have an important role in attracting and allowing for improved reproduction and development in Drosophila (Becher et al., 2012; Bellutti et al., 2018; Grangeteau et al., 2018). Nevertheless, our results indicate that the general differences we observed between fruit *versus* control media were not due to yeast content as increased performance was observed in the former media despite quite approximate yeast concentrations between media. Differences observed are likely associated with variation in carbohydrate content between fruit vs control substrates. This is because corn flour – present in the control medium but not in the fruit media, being replaced by the fruit purées – is mostly composed of a complex carbohydrate (i.e. starch) and has a very low content of simple sugars (compare P:C and P:S ratios in Table 1). This makes the carbohydrate content of the control medium not easily metabolized by Drosophila, likely reducing the efficiency of the energy metabolism cycle (Yamada et al. 2018). In contrast, the different fruits replacing corn flour in the medium have a carbohydrate content that has high levels of simple sugars (see Table 1). This likely resulted in a more efficient carbohydrate digestion potentially allowing for the recruitment of more energy metabolism resources, allocated to reproductive and developmental functions (Fanson et al. 2012).

A divergent performance across fruit hosts could have occurred has a consequence of the clearly contrasting protein to carbohydrate (P:C) ratios of the different fruits used in the experiment. This has been shown to be the case in D. suzukii, with several studies highlighting the role of P:C ratios in fecundity and development, despite some discrepancies in the direction and magnitude of the effects observed (e.g. see (Olazcuaga et al., 2019; Shu et al., 2022; Silva-Soares et al., 2017; Young et al., 2018). This possible effect was not observed in our study with *D. subobscura* as we did not detect differences in performance between fruit media. The high performance across fruits coupled with the fact that these flies were not exposed to such media during laboratory evolution suggests limited scope for differences in plasticity between fruit hosts in the founder populations. Nevertheless, it is possible that the inclusion of high yeast concentrations in all substrates – and hence high protein content - in all fruit media has contributed to reduce the impact of the distinct nutritional content of fruits particularly considering that proteins are essential for reproduction and development (Lee et al. 2008; Fanson et al. 2012). This effect can also explain in part why we did not find any evidence for “trap plants” in our study - hosts that attract Drosophila for oviposition but lack the nutritional composition that is vital for a successful development. Examples of such trap plants - helpful in mitigation strategies for pest species - have been found when analysing the performance of *D. suzukii* in different fruit hosts (Poyet et al. 2015; Olazcuaga et al. 2019).

Raspberry has been described as a high-quality fruit host for *D. suzukii* reproduction and development, with increased performance relative to other fruits (Abraham et al., 2015; Shu et al., 2022). While we found some suggestion that raspberry allowed higher average performance across fruit media for most traits studied, significant differences were only obtained when comparing raspberry to the control media for total fecundity and reproductive success. This discrepancy between species can potentially indicate that *D. suzukii* is more sensitive to varying nutritional components of the fruit substrate, although methodological differences between studies – namely in the nutritional content of the different media - might have also played a role. Such outcome is interesting considering the clear differences between these species in food habitat adaptation – *D. suzukii* mostly explores ripening fruit (e.g. (Asplen et al. 2015) whereas *D. subobscura* is more attracted to overripe, rotting fruits (Markow and O’Grady 2008) with higher levels of fermentation. In this context, important aspects that may generate differences in oviposition behaviour between species – not directly tested in our study - are the stiffness of the substrate (Karageorgi et al. 2017; Zhang et al. 2020) and the protein / yeast content (Becher et al. 2012; Silva-Soares et al. 2017), likely correlated with the ripeness of the fruit. Future studies should try to discriminate the relative importance of the different ecological / nutritional features mentioned above.

Altogether, our experiment shows that long-term lab populations of *Drosophila subobscura* (still) show increased fecundity and reproductive success when exposed to new feeding and oviposition substrates, likely more nutritive and attractive. Importantly, this occurred after long-term evolution in a particular substrate suggesting that its ability to plastically respond to diverse environmental cues was not lost in such a homogeneous environment. This argues for the utility of laboratory populations for long-term evolutionary studies (Stroud and Ratcliff 2025). Furthermore, we highlight the ability of *D. subobscura* to successfully oviposit and develop on several fruit hosts, which reinforces its potential to colonize different environments.

## Data Availability Statement

The data that supports the findings of this study are available in the supplementary material of this article (Supplementary Tables S1 and S2).

