## Supplementary Figures for "Long-term laboratory Drosophila populations prefer ancestral nutritional cues from the environment"

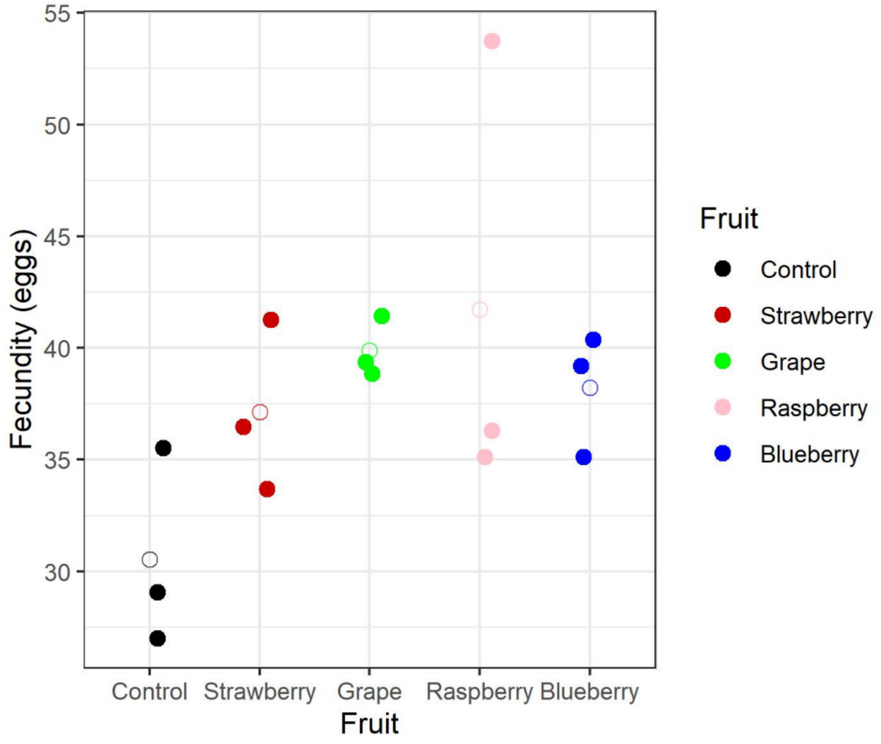


Figure S1 – Fecundity of day 7 (F7) across different media. Full circles represent each replicate population analysed in the study, while the open circles represent the average value of the three replicate populations.

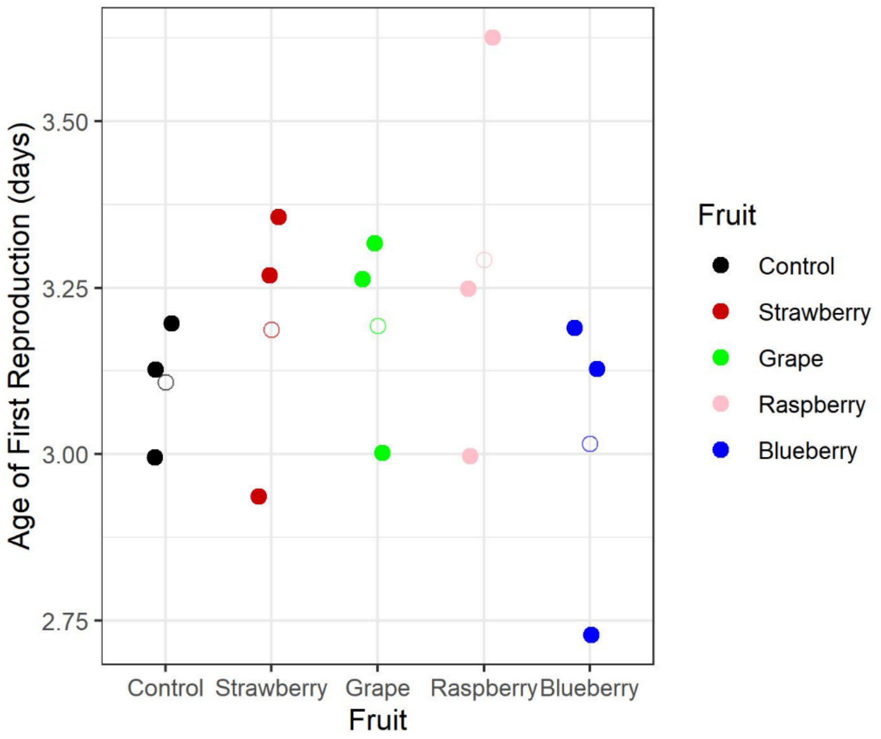


Figure S2 – Age of First Reproduction across different media. Full circles represent each replicate population analysed in the study, while the open circles represent the average value of the three replicate populations.

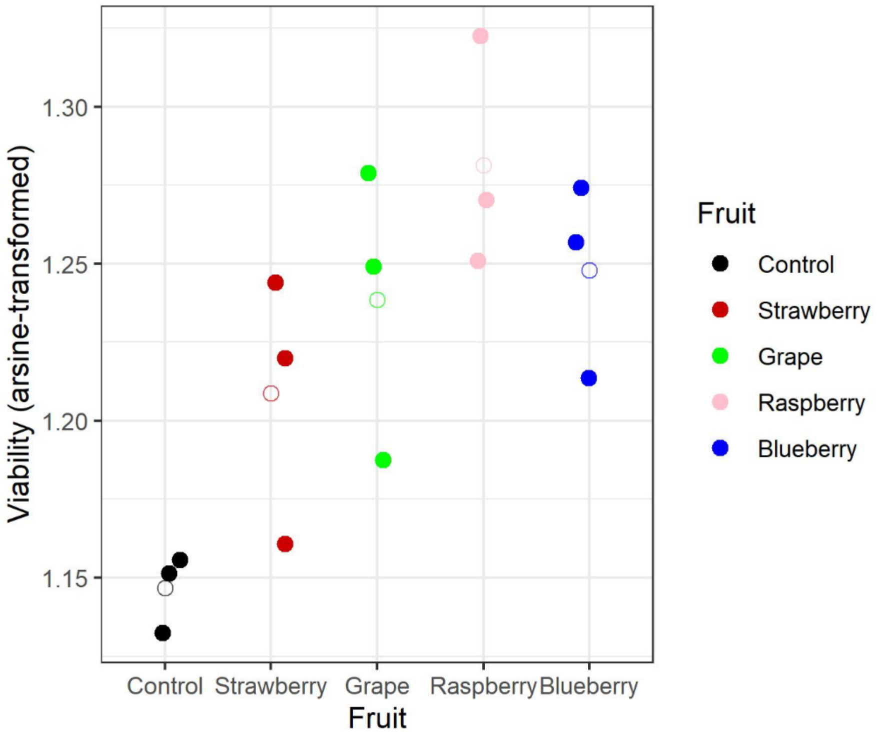


Figure S3 – Juvenile Viability across different media. Full circles represent each replicate population analysed in the study, while the open circles represent the average value of the three replicate populations.
